# NANHA - Neurostimulation in Atypical Neurodevelopment: a Harmonized Atlas

**DOI:** 10.64898/2026.09.12.751126

**Authors:** Madhumita Mondal, Camelia Guha, Sree Aswatha Suresh, Shrish Vashishth, Vignesh Muralidharan, Areejit Samal

**Affiliations:** The Institute of Mathematical Sciences (IMSc), Chennai, India; Homi Bhabha National Institute (HBNI), Mumbai, India; Department of Applied Mathematics and Computational Sciences, PSG College of Technology, Coimbatore, Tamil Nadu 641004, India; Department of Bioscience and Bioengineering, Indian Institute of Technology Jodhpur (IITJ), Jodhpur 342030, India

**Keywords:** neurodevelopmental disorder, non-invasive brain stimulation, brain region, neuromodulation, mental disorder, transcranial magnetic stimulation, transcranial electrical stimulation, database

## Abstract

Non-invasive brain stimulation (NIBS) has become an important tool for investigating brain function and exploring therapeutic intervention in neurodevelopmental disorders (NDDs). Nevertheless, relevant studies are scattered across different NDDs, stimulation modalities, target regions, and experimental designs, which makes systematic exploration and comparative analysis challenging. Therefore, we present NANHA, a manually curated, harmonized and freely accessible resource of published NIBS studies in NDDs. The database was constructed using systematic PubMed searches covering 22 NDDs and five NIBS modalities. NANHA captures participant characteristics, study design, stimulation purpose and parameters, targeted brain regions, behavioral, cognitive, and neurophysiological outcomes, safety information, and follow-up assessments, whenever available. The NANHA web platform (https://cb.imsc.res.in/nanha/) provides searchable and downloadable records, advanced filtering options, and interactive visualizations to support data exploration. Overall, NANHA is a standardized resource for evidence synthesis, comparative analyses, identification of research gaps, and the development of disease-specific stimulation protocols for NDDs.

## Background and Summary

Neurodevelopmental disorders (NDDs) comprise a heterogeneous group of conditions characterized by atypical brain development that affects cognitive, behavioral, motor, language, and social functioning. Accumulating evidence suggests that many NDDs are associated with disruptions in the finely regulated balance between cortical excitation and inhibition (E/I balance), which leads to altered neural circuit development and function^1–4^. Consequently, modulating excitation/inhibition (E/I) dynamics has emerged as an important therapeutic strategy for targeting improvements in several NDDs.

Non-invasive brain stimulation (NIBS) has therefore become an important non-pharmacological approach for both probing cortical physiology and modulating atypical neural activity^5^. Techniques such as transcranial magnetic stimulation (TMS)^6^, transcranial electrical stimulation (tES)^7^, including transcranial direct current stimulation (tDCS)^8^, transcranial alternating current stimulation (tACS)^9^, transcranial random noise stimulation (tRNS)^10^, and acoustic methods such as transcranial focused ultrasound stimulation (tFUS)^11^ are increasingly being applied to investigate cortical excitability and plasticity to evaluate their therapeutic potential across different NDDs.

The clinical adoption of these techniques has been rapid, however, the literature around the effects of NIBS on NDDs remain largely fragmented, spread across different stimulation modalities, cortical targets and outcome measures^12–15^. For instance, the effects of NIBS across a range of NDDs have been investigated to evaluate its influence on motor, cognitive, behavioral, and neurophysiological outcomes^16–18^. These studies have also explored diverse stimulation targets, spanning several cortical sites and protocols (modulating intensity and stimulation frequency) to understand disease mechanisms better and assess the therapeutic potential of NIBS. A recent systematic review summarized randomized controlled trials of selected NIBS modalities in a limited number of NDDs^19^. Crucially, no open-access or publicly available, literature-curated standardized database exists that integrates participant baseline characteristics, stimulation parameters, study design, and reported outcomes across multiple NIBS modalities and NDDs. Therefore, to address this gap, we developed Neurostimulation in Atypical Neurodevelopment: a Harmonized Atlas or NANHA (https://cb.imsc.res.in/nanha/), a meticulously curated and standardized database at the intersection of NIBS and NDDs that provides a comprehensive, searchable resource for exploring the current evidence of published NIBS studies across different NDDs. It includes multiple neuromodulation modalities: TMS, tDCS, tACS, tRNS, and tFUS. NANHA integrates participant demographics (e.g., sample size, mean or median age, age range, and sex), clinical characteristics (e.g., intellectual ability, comorbidities, and medication status), study design, stimulation protocols, targeted brain regions, and behavioral, cognitive, and neurophysiological outcomes, together with safety, follow-up, and long-term outcome information, whenever available. By harmonizing study-level metadata across stimulation paradigms, the resource allows systematic evidence synthesis, cross-study comparisons, identification of research gaps, and investigation of the underlying circuit biotypes governing neuromodulation responses across traditional diagnostic boundaries.

## Methods

### Compilation and curation of data on application of NIBS in individuals with NDDs

The NDDs included in the database were primarily based on the classification proposed by Bishop^20^, which comprises a total of 35 NDDs. In addition, we included communication disorders (CD) to capture disorders such as childhood-onset fluency disorder (stuttering). CD is classified under the Neurodevelopmental Disorders chapter of the Diagnostic and Statistical Manual of Mental Disorders, Fifth Edition (DSM-5)^21^. Therefore, a total of 36 NDDs were considered in this study. For NIBS, we focused on seven commonly used modalities: TMS, tDCS, tACS, tRNS, tFUS, transcranial interference stimulation (tIS), and transcutaneous peripheral nerve stimulation (tPNS).

To identify relevant studies, a systematic literature search was conducted using PubMed (https://pubmed.ncbi.nlm.nih.gov/), which served as the primary bibliographic database. For each of the 36 NDDs, a separate search query was performed by combining the disorder name, along with its commonly used synonyms and abbreviations, and with a comprehensive set of keywords representing the NIBS modalities considered in this study. This literature search was last conducted on 12 April 2026. As an example, one of the search queries that targeted studies on ADHD is the following:

> *(ADHD OR “attention deficit hyperactivity disorder” OR “attention deficit/hyperactivity disorder”) AND (NIBS OR “noninvasive brain stimulation” OR TMS OR “transcranial magnetic stimulation” OR tDCS OR “transcranial direct current stimulation” OR tACS OR “transcranial alternating current stimulation” OR tRNS OR “transcranial random noise stimulation” OR tFUS OR TUS OR “transcranial focused ultrasound” OR tIS OR “transcranial interference stimulation” OR tPNS OR “transcutaneous peripheral nerve stimulation”)*

The complete search queries for each NDD, along with their abbreviations and the number of articles retrieved, are provided in **Table S1**. This systematic search formed the basis for the subsequent screening and curation of studies included in the database. The PubMed search identified a total of 1,942 papers across 26 of the 36 NDDs included in this study, whereas no articles were available for the other 10 disorders (see **Table S1**).

Next, we followed a three-stage procedure to refine the set of articles retrieved from the literature search. First, duplicate records were removed from the results of the 26 separate search queries, yielding 1,726 unique articles. Second, the retrieved articles were screened based on their titles and abstracts for relevance. The inclusion criteria were: (1) studies involving individuals with one of the selected NDDs, (2) studies employing one or more of the seven selected NIBS modalities, (3) peer-reviewed original research articles, and (4) studies reporting sufficient demographic information of the participants, methodological details on the stimulation protocol, and target brain regions. The exclusion criteria included review articles, meta-analyses, non-English publications, study protocols, duplicate records, studies not related to NDDs or NIBS, and articles without accessible full text. This screening stage excluded 912 articles and the remaining 814 relevant articles were retained for further evaluation. Third, the full texts of the screened articles were assessed to confirm their eligibility according to the same inclusion and exclusion criteria. Studies that did not provide the required information for data extraction were excluded. Following this detailed assessment, data were extracted from 526 eligible articles and incorporated into the final dataset. The overall literature search and study selection process is summarized in a PRISMA flow diagram (**Figure 2**).

**Figure 1.**
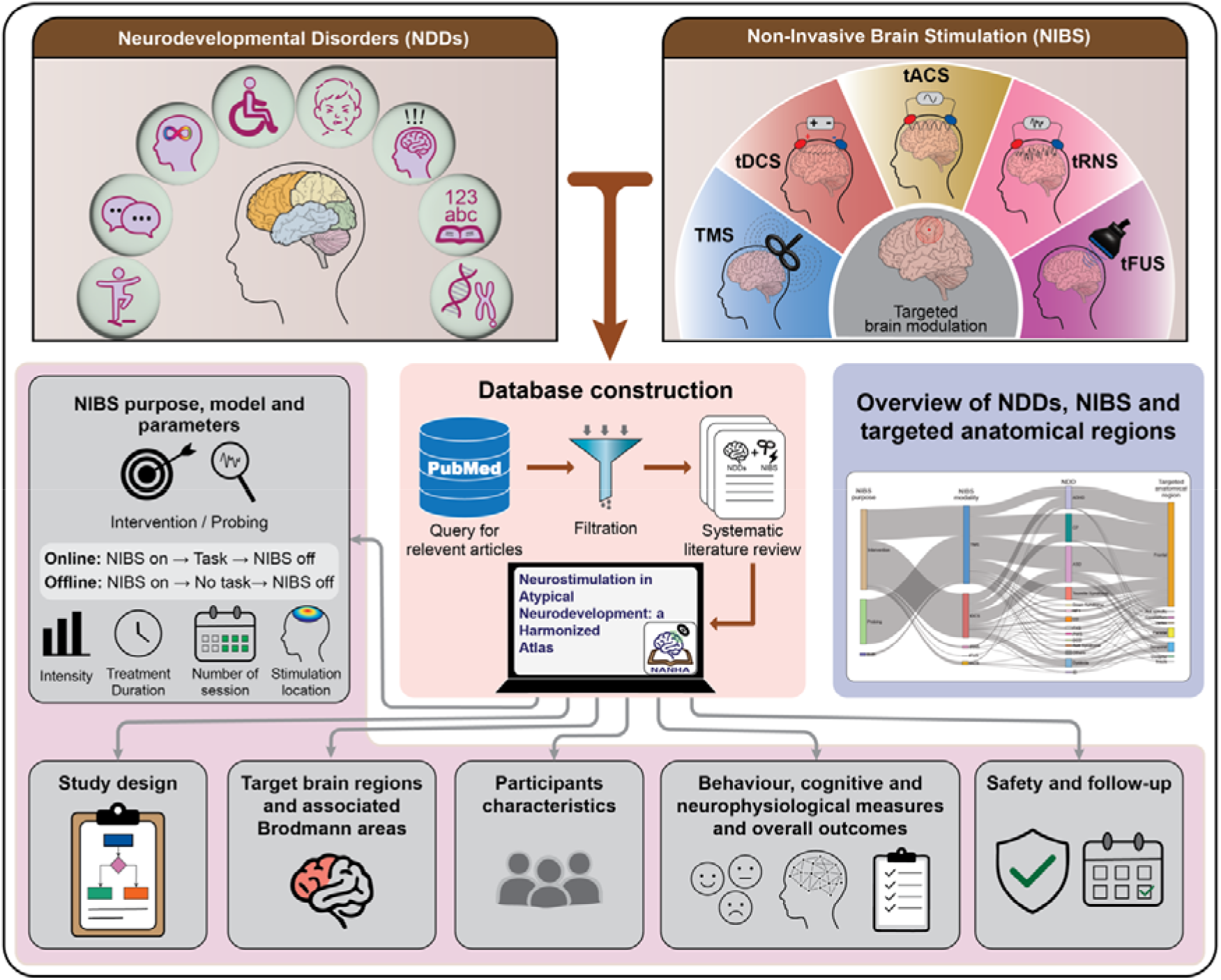
Schematic workflow illustrating the compilation and curation of neurodevelopmental disorder (NDD) and non-invasive brain stimulation (NIBS) literature, extraction of study-level information and parameters, and their representation in the NANHA web interface.

**Figure 2.**
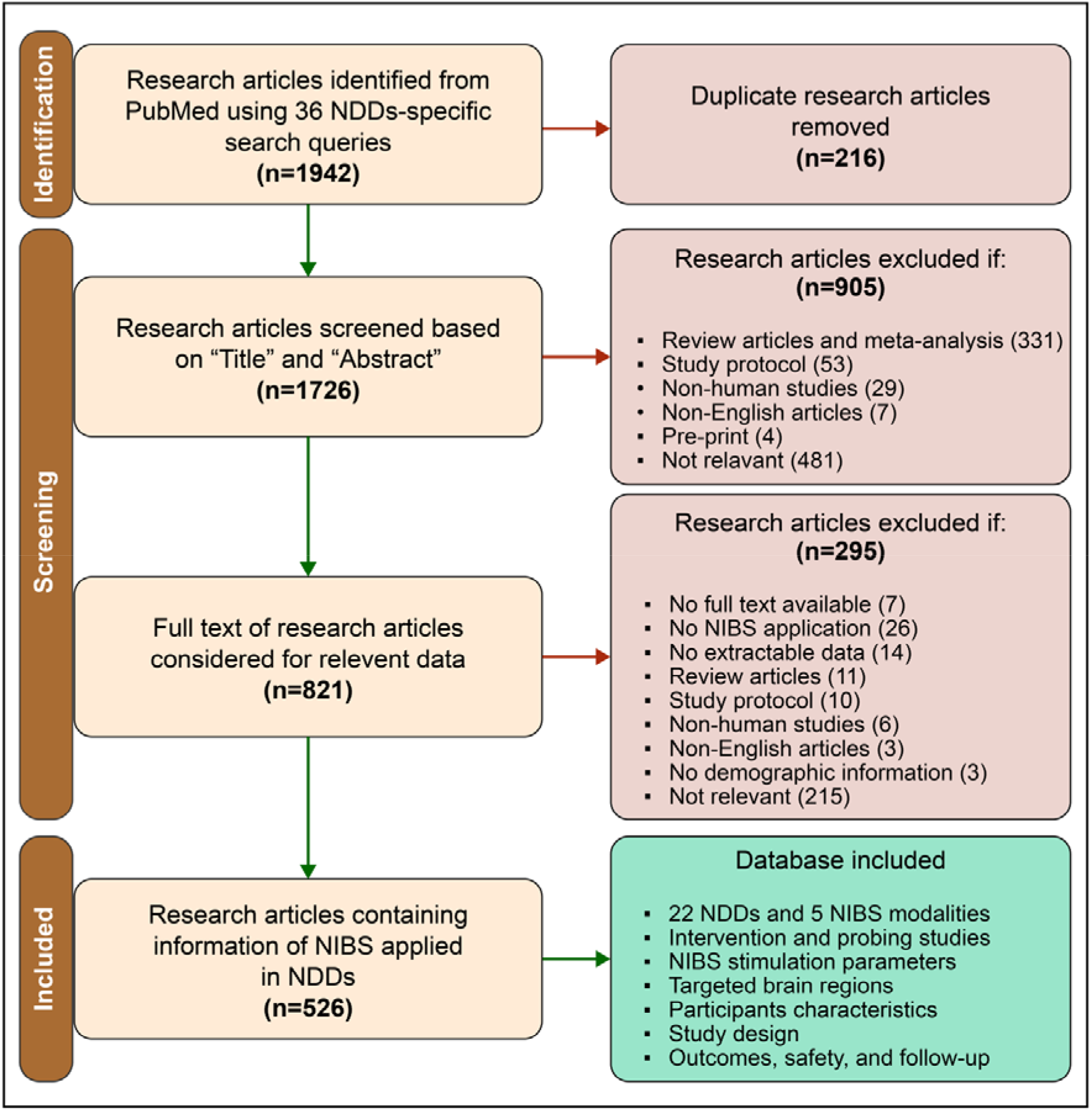
A PRISMA flow diagram summarizing the literature search, screening, and study selection process used to identify publications reporting the application of non-invasive brain stimulation (NIBS) in neurodevelopmental disorders (NDDs).

Next, data were manually extracted from the 526 articles, covering 22 NDDs and five NIBS modalities (TMS, tDCS, tACS, tRNS, and tFUS). Extracted variables included the type of NDD in the experimental group, control condition, participant characteristics (e.g., mean age, sex, IQ, and comorbid conditions), study design, target brain region, stimulation modality (e.g., TMS, tDCS, and tACS), stimulation purpose (intervention or probing), stimulation model (online or offline), stimulation protocol (including frequency, intensity, motor threshold, pulse number, neuronavigation, electrode positions, session duration, and number of sessions), primary outcome measures (behavioral, cognitive, and neurophysiological), safety outcomes, and long-term effects. Stimulation model was categorized as online or offline based on its timing relative to the behavioral or cognitive task. Online stimulation refers to NIBS delivered during the task, whereas offline stimulation refers to NIBS delivered before or after the task. Additionally, some of the 526 eligible studies have carried out interventions other than NIBS; in such studies, mostly NIBS, particularly TMS, has been used to probe brain activity.

### Data standardization and annotation

Each included study was initially represented by a single record. However, when a study contained multiple independent experimental conditions, participant groups, control groups, stimulation targets, or NIBS modalities, separate records were created for each unique comparison or stimulation condition. For example, in studies including anodal, cathodal, and sham stimulation, two records were generated: anodal versus sham and cathodal versus. sham. Likewise, if different brain regions were stimulated in separate sessions (e.g., 4 target areas), each target-specific condition was recorded as an individual entry. All such records were linked to the same PubMed identifier (PMID) to preserve their common study origin while allowing condition-specific extraction of stimulation parameters, participant characteristics, and outcome measures. This approach ensured that each row represented a single, well-defined experimental comparison, which enabled accurate analysis of stimulation-specific effects without conflating results across different conditions within the same study. Any inconsistencies in terminology across studies were carefully harmonized. To facilitate cross-study comparisons, stimulation targets were subsequently classified into broader neuroanatomical regions such as: frontal, temporal, parietal, occipital, cerebellar, and insular. The distributions of the included studies across the 22 NDDs, five NIBS modalities, NIBS purpose, and neuroanatomical regions are presented in **Figure 3**. Note that, studies reporting multiple NDDs, NIBS modalities, or neuroanatomical regions contributed separately to each category, resulting in a total of 598 entries.

**Figure 3.**
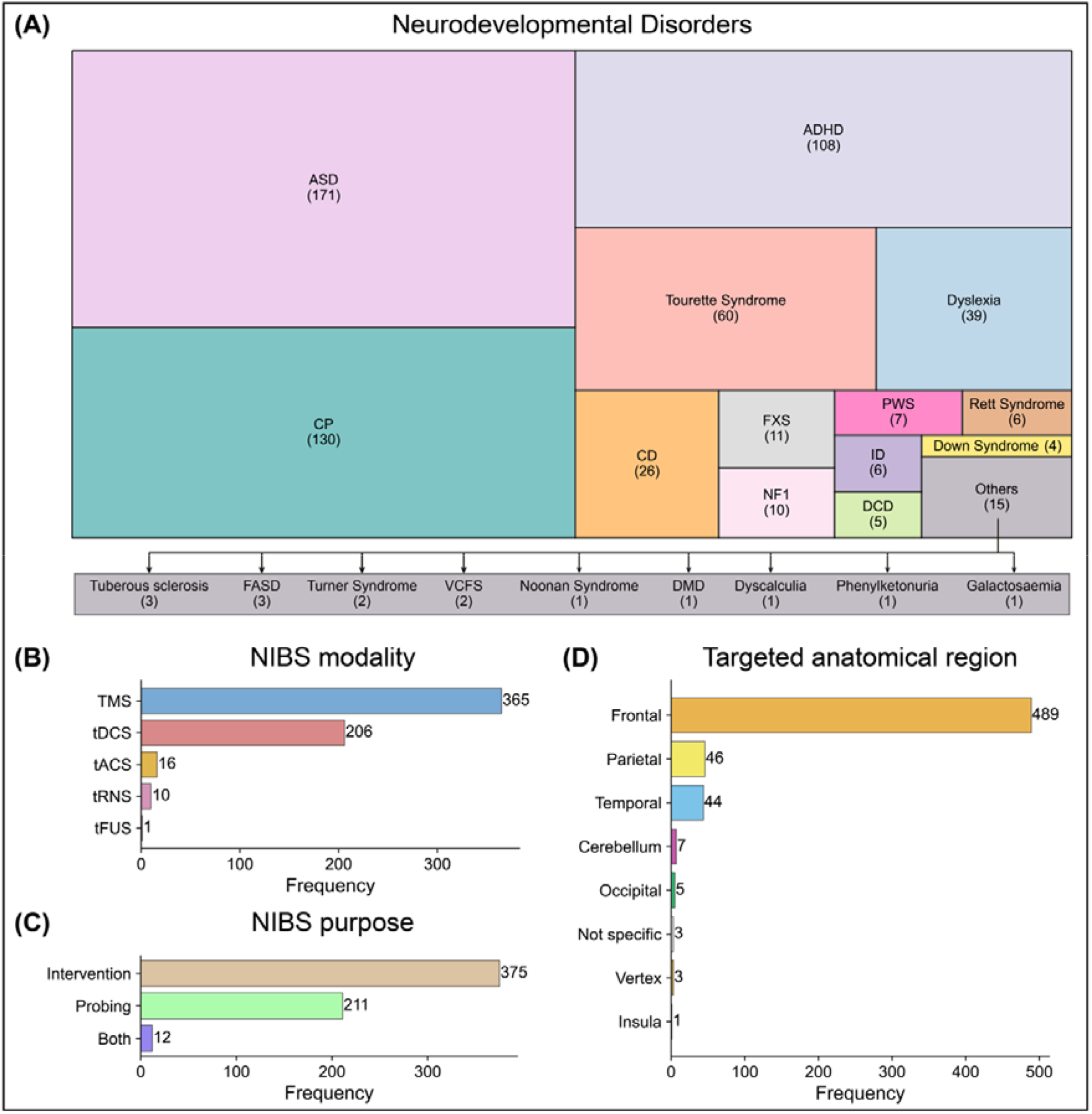
Distribution of studies included in NANHA according to **(A)** 22 neurodevelopmental disorders, **(B)** five non-invasive brain stimulation (NIBS) modalities, **(C)** three stimulation purposes (intervention, probing, or both), and **(D)** eight targeted anatomical brain regions. The full forms of all abbreviations used in the figure are as follows: ADHD: attention-deficit/hyperactivity disorder, ASD: autism spectrum disorder, CP: cerebral palsy, CD: communication disorder, DCD: developmental coordination disorder, DMD: Duchenne muscular dystrophy, FASD: fetal alcohol spectrum disorder, FXS: fragile X syndrome, ID: intellectual disability, NF1: neurofibromatosis type 1, PWS: Prader-Willi syndrome, VCFS: velocardiofacial syndrome, TMS: transcranial magnetic stimulation, tDCS: transcranial direct current stimulation, tACS: transcranial alternating current stimulation, tRNS: transcranial random noise stimulation, tFUS: transcranial focused ultrasound stimulation.

### Implementation of the NANHA web interface and database

The curated resource is publicly available through the NANHA web platform, wherein the curated dataset is downloadable in standard formats without user registration for non-commercial use. The resource is organized into separate sections for intervention and probing, to simplify literature exploration. Within each section, searches can be made by selecting one or more NDDs together with a specific NIBS modality. For standardized classification, the NDDs included in this database are linked to their corresponding International Classification of Diseases, 11th Revision (ICD-11) codes^22^. The search can be further refined using optional filters for stimulation target, stimulation model (online or offline), whether the study is a case study, and the inclusion of a control group. In addition to disorder-based searches, users can also filter out the studies by targeted brain regions across different NDDs. A dedicated help section with usage instructions and video tutorials is also included to support efficient navigation and data retrieval. All content in NANHA is available under the Creative Commons Attribution-NonCommercial-NoDerivatives 4.0 International (CC BY-NC-ND 4.0) license (https://creativecommons.org/licenses/by-nc-nd/4.0/).

All curated records are maintained in a MariaDB (https://mariadb.org/) database and accessed via Structured Query Language (SQL) queries. The web interface was developed using HTML5, CSS3, JavaScript, and PHP (https://www.php.net/) and is hosted on an Apache HTTP Server (https://httpd.apache.org/) running on Debian 9.13. Interactive 3D brain visualizations were implemented using Plotly.js (https://plotly.com/javascript/), and MNE-Python (https://mne.tools/stable/index.html), FreeSurfer (https://surfer.nmr.mgh.harvard.edu/), and NiBabel (https://pypi.org/project/nibabel/) used for surface mesh generation and processing. FreeSurfer fsaverage, the PALS_B12 Brodmann Parcellation Atlas, and the Talairach Daemon (https://talairach.org/) were used for cortical surface representation, Brodmann area annotation, and anatomical coordinate mapping, respectively. Statistical charts and data distributions presented on the platform were developed using Chart.js JavaScript library (https://www.chartjs.org/).

#### Data records

In the curated dataset, the relationships amongst the NIBS purpose, NIBS modality, NDDs, and targeted anatomical regions are summarized in the Sankey diagram (**Figure 4**). For interactive exploration, the curated dataset is publicly available through the NANHA web interface at https://cb.imsc.res.in/nanha/.

**Figure 4.**
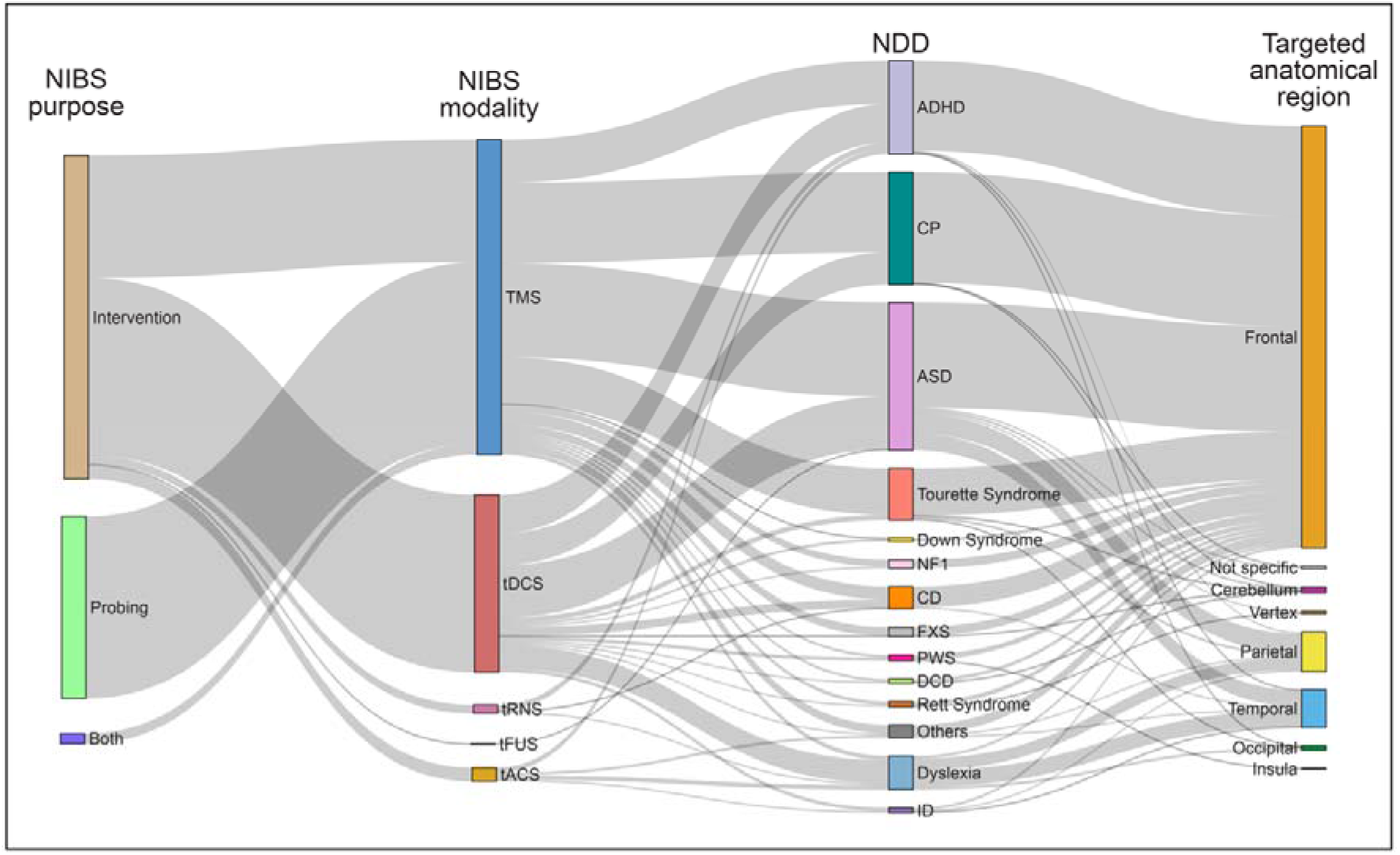
Sankey diagram illustrating the relationships between stimulation purpose, non-invasive brain stimulation (NIBS) modality, neurodevelopmental disorders (NDDs), and targeted anatomical regions. The width of each flow is proportional to the number of studies sharing the corresponding combination. The full forms of all abbreviations used in the figure are as follows: TMS: transcranial magnetic stimulation, tDCS: transcranial direct current stimulation, tACS: transcranial alternating current stimulation, tRNS: transcranial random noise stimulation, tFUS: transcranial focused ultrasound stimulation, ADHD: attention-deficit/hyperactivity disorder, ASD: autism spectrum disorder, CP: cerebral palsy, CD: communication disorder, DCD: developmental coordination disorder, FXS: fragile X syndrome, ID: intellectual disability, NF1: neurofibromatosis type 1, PWS: Prader-Willi syndrome.

### Study-Level Metadata

This datasheet (**Table S2**) with 798 rows and 49 columns contains the primary curated information extracted from each study included in the dataset. Each row corresponds to a unique experimental condition or comparison within a study and is linked to its original publication through the PMID. The datasheet includes the study identifiers, NDD, and participant characteristics, including sample size, age (mean, median, and range where available), sex, intelligence quotient (IQ) score, comorbid conditions, medication status, and control group information. Study-level information, including the study design (e.g., parallel, cross-over, case-control, single-arm, or cross-sectional), randomization, blinding, NIBS modality and purpose and model, stimulation target, associated Brodmann areas (BAs) and brain lobe, is also provided. Additionally, the database summarizes the behavioral and cognitive tasks performed, along with the overall behavioral and cognitive outcomes reported in each study. When multiple outcome measures were assessed, the reported outcome reflects the overall study findings. Studies with both positive and negative changes across different measures were classified as mixed; otherwise, the reported direction of change (improved, reduced, or not changed) was mentioned. Neurophysiological changes, mainly reported in TMS studies, were extracted and categorized based on their overall effect (e.g., increase, decrease, modulated, or no change). The extracted measures include motor evoked potential (MEP) amplitude, intracortical inhibition (ICI), intracortical facilitation (ICF), resting motor threshold (RMT), active motor threshold (AMT), and other reported neurophysiological measures in the studies. Finally, information on participant dropout, adverse effects, long-term outcomes, and follow-up assessments is provided where available. When a single publication reported multiple experimental conditions, stimulation targets, or intervention modalities, separate rows were created for each condition to enable condition-specific analyses while preserving the association with the original study.

### Stimulation protocols

To summarize the stimulation protocols used in each of the five NIBS modalities: TMS, tDCS, tACS, tRNS, or tFUS, we created five datasheets (**Tables S3-S7**) corresponding to each of the NIBS modalities. Each row represents a unique experimental comparison and includes the associated PMID, study number, NDD, stimulation purpose, stimulation model, target brain region, the number and frequency of treatment sessions, treatment duration, and modality-specific stimulation parameters. These parameters are described below for each of the five NIBS modalities.

1. **TMS Parameters:** This datasheet (**Table S3**) with 508 rows and 14 columns, summarizes the stimulation protocols used in TMS studies. The TMS parameters include stimulation frequency, motor threshold, total pulses delivered per session, inter-train interval, and the use of neuronavigation.
2. **tDCS Parameters:** This datasheet (**Table S4**) with 258 rows and 12 columns, summarizes the stimulation protocols used in tDCS studies. The tDCS parameters include current intensity and the locations of the anodal and cathodal electrodes.
3. **tACS Parameters:** This datasheet (**Table S5**) with 19 rows and 13 columns, summarizes the stimulation protocols used in tACS studies. The tACS parameters include current intensity, frequency of stimulation, and the locations of the two stimulation electrodes.
4. **tRNS Parameters:** This datasheet (**Table S6**) with 12 rows and 12 columns, summarizes the stimulation protocols used in tRNS studies. The tRNS parameters include stimulation intensity, frequency of stimulation, and the locations of the stimulation electrodes.
5. **tFUS Parameters:** This datasheet (**Table S7**) with one row and 13 columns, summarizes the stimulation protocols used in tFUS studies. The tFUS parameters include ultrasound frequency, acoustic intensity (spatial-peak temporal-average intensity (ISPTA) and spatial-peak pulse-average intensity (ISPPA)), pulse repetition frequency (PRF), and duty cycle.

### Abbreviation

This datasheet (**Table S8**) contains all the abbreviations used in the database.

#### Technical validation

To ensure the accuracy, consistency, and reliability of the database, a multi-stage validation process was carried out throughout data collection and curation. All studies underwent manual screening based on predefined inclusion and exclusion criteria. Only original human studies reporting NIBS experiments in individuals with NDDs were included in the final database.

Data from all included studies were extracted using a standardized template to reduce variability and ensure a consistent representation of studies. Study characteristics, participant demographics, stimulation parameters, outcome measures, and safety-related information were initially extracted from each study. In a subsequent step, all data were harmonized using standardized terminology. For instance, when the mean and standard error (SE) of participants’ age were reported, the SE was converted to a standard deviation (SD) to achieve consistency across datasets. Whenever possible, reported values were recorded directly from the original publication without modification.

However, target brain regions were reported inconsistently across studies, using anatomical names, BAs, EEG electrode positions, or functional region names. To ensure consistency, all reported stimulation targets were manually standardized and annotated with the corresponding BAs and subsequently mapped into broader neuroanatomical regions (e.g., frontal, parietal, temporal, occipital, insular, cerebellar, and vertex)^23–27^. This helps to enable consistent cross-study comparisons and identify the brain lobes most commonly targeted across different NDDs. Notably, the BA annotation was restricted to cortical areas corresponding to NIBS-targetable surface regions; medial and deeper cortical areas were not included in the BA mapping.

#### Usage notes

The NANHA resource is intended to serve as a comprehensive resource for the systematic exploration and comparison of NIBS studies across different NDDs. Some of the potential applications of the resource are listed below.

### Transdiagnostic assessment of NIBS treatment effects

Users can quantify the effectiveness of different NIBS modalities across NDDs by assessing changes in outcome measures from baseline and determining whether these changes differ from those observed in the control group. The control group can be both the neurotypical population or a stimulation control, for instance sham. This provides a framework for comparing treatment effects across disorders and stimulation modalities, and for exploring common patterns of treatment response and sources of heterogeneity across studies. This also allows for identifying sources of variance in the stimulation effects for different NDDs. For example, by holding the protocol constant (say, 10Hz rTMS), users can look at the outcome measures across ADHD, Autism and Dyslexia.

### Mapping circuit plasticity and spatial target engagement biomarkers

The dataset will allow the users to investigate studies in which NIBS is used as a probing modality or as an intervention followed by probing to characterize stimulation-induced neurophysiological changes, especially cortical excitation-inhibition changes. Specifically, users can understand the relationship between the stimulated brain region and alterations in cortical excitability and inhibition, quantified using measures such as motor evoked potential (MEP) amplitude, short-interval intracortical inhibition (SICI), long-interval intracortical inhibition (LICI), cortical silent period (CSP), intracortical facilitation (ICF), resting motor threshold (RMT), and active motor threshold (AMT). This analysis enables systematic mapping of circuit plasticity, identifies region-specific patterns of target engagement, and provides insights into how different cortical targets modulate excitatory and inhibitory neural circuits.

### Network-level evaluation of stimulation modalities, target regions, and protocol transferability

The database provides harmonized information on stimulation modalities, cortical target regions, stimulation protocols, and associated behavioral outcomes across different NDDs. These data can be used to compare the effectiveness of different NIBS modalities and target regions through systematic reviews. Furthermore, they enable evaluation of whether identical stimulation protocols produce comparable effects across different cortical targets and thereby provide insights into the spatial specificity and transferability of NIBS protocols.

### Quantifying safety and tolerability to stimulation modalities

The database provides information on adverse events, stimulation modalities, and stimulation parameters, including intensity and frequency, reported across studies. These data can be used to systematically evaluate the safety and tolerability of different NIBS protocols and investigate associations between stimulation parameters and the occurrence of adverse events. Such analyses may help identify parameter ranges associated with favorable safety profiles and inform the design of future stimulation protocols, particularly those employing higher stimulation intensities or more focal and penetrating stimulation approaches.

### Unravelling inter-individual response variability in tES

tES (tDCS/tACS/tRNS) is characterized by substantial inter-individual variability, where identical stimulation parameters often lead to different outcomes across participants. These data can be used to investigate factors contributing to treatment response variability by examining participant characteristics, including medication use, comorbid conditions, intellectual ability, age, and control conditions. Such analyses may improve the understanding of tES efficacy and enhance comparisons of treatment effects relative to sham or neurotypical groups.

### Shift towards neurotypicals based on stimulation and NDD type

This dataset can also be used to investigate which kinds of neurostimulation are most effective in improving clinical, behavior, or cognitive outcomes toward those observed in neurotypicals. This will provide an evidence-based approach for developing and refining disease-specific stimulation protocols and guide the design of future therapeutic interventions for different neurodevelopmental disorders.

## Data availability

The curated dataset is publicly accessible through the NANHA web interface (https://cb.imsc.res.in/nanha/).

## Author contributions

*Conceptualization:* Madhumita Mondal, Vignesh Muralidharan, Areejit Samal; *Methodology:* Madhumita Mondal, Camelia Guha, Sree Aswatha Suresh, Shrish Vashishth, Vignesh Muralidharan, Areejit Samal; *Data curation:* Madhumita Mondal, Camelia Guha, Shrish Vashishth; *Formal analysis:* Madhumita Mondal, Camelia Guha, Vignesh Muralidharan, Areejit Samal; *Visualization:* Madhumita Mondal, Sree Aswatha Suresh, Vignesh Muralidharan, Areejit Samal; *Software:* Madhumita Mondal, Camelia Guha, Sree Aswatha Suresh, Vignesh Muralidharan, Areejit Samal; *Supervision:* Areejit Samal; *Writing – original draft:* Madhumita Mondal, Vignesh Muralidharan, Areejit Samal; *Writing – review & editing:* Madhumita Mondal, Vignesh Muralidharan, Areejit Samal.

## Supplementary Tables

**Table S1. List of considered neurodevelopmental disorders (NDDs), their abbreviations, PubMed search queries, and the number of articles retrieved for each NDD.** The table lists the NDDs considered in the literature search, their corresponding abbreviations, search terms used in PubMed, and the number of articles retrieved for each NDD. The PubMed search strategy consisted of two components: NDD-related search terms and non-invasive brain stimulation (NIBS)-related search terms, which were combined using the AND operator for each disorder. The PubMed search was last carried out on 12 April 2026.

**Table S2. This table contains the primary curated information extracted from 526 studies.** Each row corresponds to a unique experimental condition or comparison within a study and is linked to its original publication through the PMID. The table includes the unique study identifiers, neurodevelopmental disorder (NDD), participant characteristics, including sample size, age (mean, median, and range where available), sex, intelligence quotient (IQ) score, comorbid conditions, medication status, control group information, study-level information, including the study design (e.g., parallel, cross-over, case-control, single-arm, or cross-sectional), randomization, blinding, NIBS modality, purpose, and model, stimulation target, associated Brodmann areas and brain lobe, behavioral and cognitive tasks performed, along with the overall behavioral and cognitive outcomes reported in each study, neurophysiological changes (e.g., increase, decrease, modulated, or no change). The extracted neurophysiological measures include motor evoked potential (MEP) amplitude, intracortical inhibition (ICI), intracortical facilitation (ICF), resting motor threshold (RMT), active motor threshold (AMT), and other reported neurophysiological measures in the studies. Finally, information on participant dropout, adverse effects, long-term outcomes, and follow-up assessments is provided where available. When a single publication reported multiple experimental conditions, stimulation targets, or intervention modalities, separate rows were created for each condition to enable condition-specific analyses while preserving the association with the original study.

**Table S3: Transcranial magnetic stimulation (TMS) parameters**. This table summarizes the stimulation protocols and parameters used in TMS studies. Each row represents a unique experimental comparison and includes the associated unique study identifier, neurodevelopmental disorder (NDD), stimulation purpose, stimulation model, target brain region, the number and frequency of treatment sessions, treatment duration, TMS modality, stimulation frequency, motor threshold, total pulses delivered per session, inter-train interval, and the use of neuronavigation.

**Table S4: Transcranial direct current stimulation (tDCS) parameters**. This table summarizes the stimulation protocols and parameters used in tDCS studies. Each row represents a unique experimental comparison and includes associated unique study identifier, neurodevelopmental disorder (NDD), stimulation purpose, stimulation model, target brain region, the number and frequency of treatment sessions, treatment duration, tDCS modality, current intensity, and the locations of the anodal and cathodal electrodes.

**Table S5: Transcranial alternating current stimulation (tACS) parameters**. This table summarizes the stimulation protocols and parameters used in tACS studies. Each row represents a unique experimental comparison and includes associated unique study identifier, neurodevelopmental disorder (NDD), stimulation purpose, stimulation model, target brain region, the number and frequency of treatment sessions, treatment duration, tACS modality, current intensity, frequency of stimulation, and the locations of the two stimulation electrodes.

**Table S6: Transcranial random noise stimulation (tRNS) parameters**. This table summarizes the stimulation protocols and parameters used in tRNS studies. Each row represents a unique experimental comparison and includes associated unique study identifier, neurodevelopmental disorder (NDD), stimulation purpose, stimulation model, target brain region, the number and frequency of treatment sessions, treatment duration, tRNS modality, stimulation intensity, frequency of stimulation, and the locations of the stimulation electrodes.

**Table S7: Transcranial focused ultrasound stimulation (tFUS) parameters**. This table summarizes the stimulation protocols and parameters used in tFUS studies. Each row represents a unique experimental comparison and includes associated unique study identifier, neurodevelopmental disorder (NDD), stimulation purpose, stimulation model, target brain region, the number and frequency of treatment sessions, treatment duration, tFUS modality, ultrasound frequency, Acoustic intensity (spatial-peak temporal-average intensity (ISPTA) and spatial-peak pulse-average intensity (ISPPA)), pulse repetition frequency (PRF), and duty cycle.

**Table S8: Abbreviations**. This table contains all the abbreviations and their full forms used across the NANHA database.

